# Biologically Informative NA Deconvolution (BIND) excavates hidden features of the proteome from missing values in large-scale datasets

**DOI:** 10.1101/2025.06.19.660508

**Authors:** Guo Weiheng, Jin Wenyi, Zheng Jieyi, Pan Yilin, Wang Rui, Zhang Jian, Feng Xikang, Chen Lingxi, Zhang Liang

## Abstract

The fast-advancing mass spectrometry and related technologies have greatly extended the depth of coverage in large-scale proteomics studies, including single-cell applications. As sample numbers grow rapidly, it is often challenging to interpret the proteins with missing values that are often presented as “NA” (not available). It could be the evidence of no expression, low expression below the detection threshold, or false negative detection due to technical issues. Existing methods for missing values imputation, while generally useful, rarely consider the non-random NA values that inform biological significance. In the current study, we developed **B**iologically **I**nformative **N**A **D**econvolution (BIND) that applies an adaptive neighborhood-based modeling to deconvolve the nature of NAs as “biological” (low/no expression) or technical (experimental errors). Applying to multiple cell line datasets and human tissue extracellular vesicle datasets, BIND excavated the NAs that indicated “hallmark absence” of unique proteins. This led to improvements in protein-protein interaction analysis and the identification of novel disease biomarkers. To facilitate its public accessibility, we compiled BIND into a web server that features functional online operations and interactive visualizations. Furthermore, we demonstrated that the BIND server could deconvolve the NAs and improve the analyses of single-cell proteomics datasets. Overall, BIND delineates the biological significance of missing values rather than treating them as a burden, providing a critical perspective for understanding the complex proteome in various biological contexts.

**Graphical abstract:** 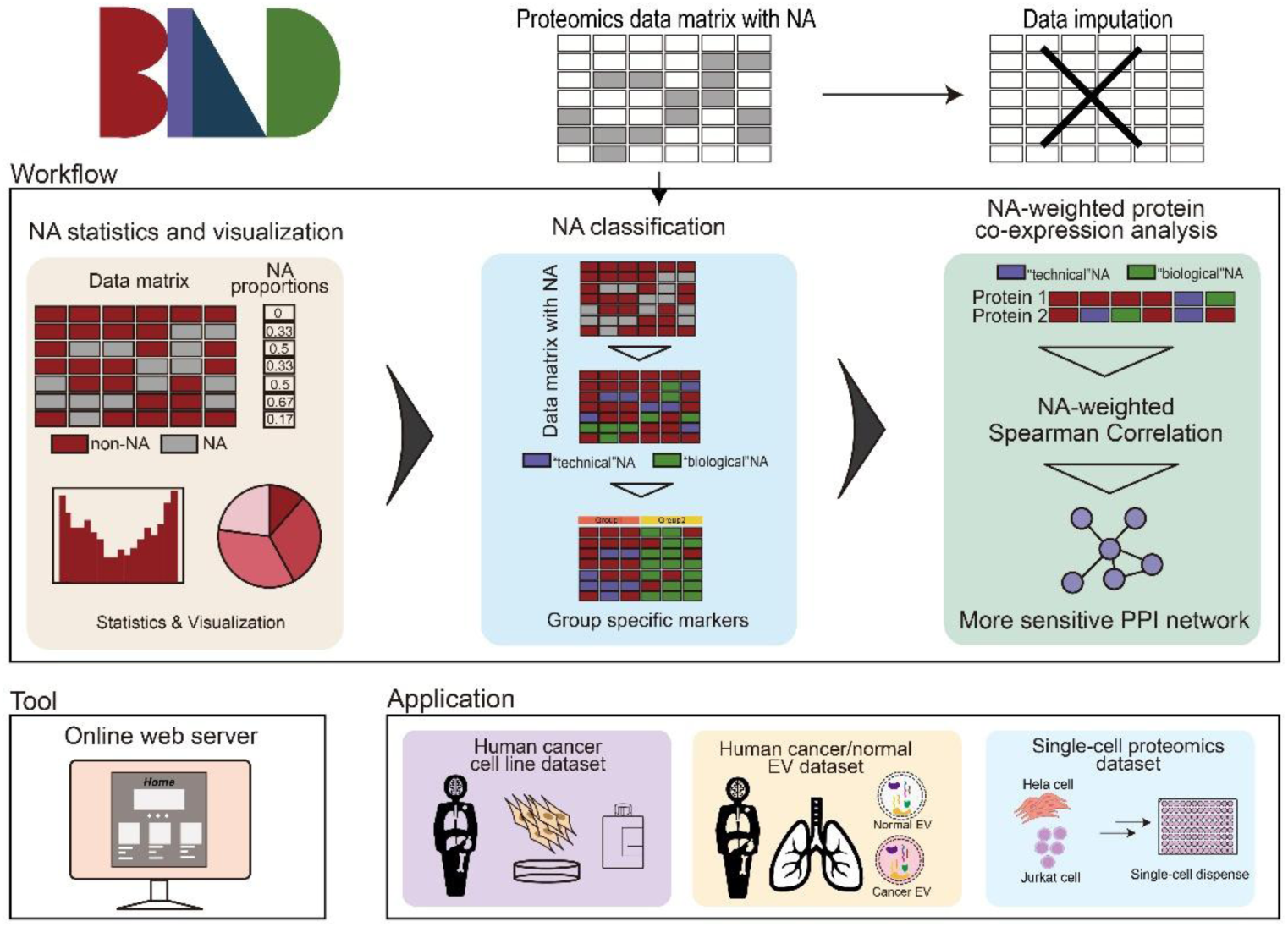

## Introduction

Proteomics has become integral to fields including biology, medicine, and environmental science^1^. However, limitations in sample processing and instrument detection often lead to missing values in proteomics datasets^2^. The missing values are often presented as “NA” (not available), which complicates the subsequent extraction of biological insights from the datasets. For example, statistical analyses like principal component analysis and hierarchical clustering are difficult to execute with an incomplete data matrix. Although advancements in mass spectrometry technology have extended the depth of protein identification and quantification, missing values remain a challenging issue and even get exacerbated as sample numbers increase^3^.

Missing values in proteomics are often categorized as missing completely at random (MCAR), missing at random (MAR), and missing not at random (MNAR)^4^. MCAR refers to data missing due to multiple small errors and random fluctuations, which are independent of the protein abundance. MAR considers conditional dependencies and is generally grouped with MCAR in proteomics. These two types of missing values are usually due to technical considerations. In contrast, MNAR is typically due to biological factors. It correlates with protein and peptide abundance, with a higher likelihood of missing for low abundances that are close to the instrumental detection limit. The distribution typically shows left-censored deletions^5–7^. Decoding the nature of missing values is crucial for trustworthy analysis and decision-making in modern proteomics.

To address the challenges brought by missing values, data imputation is a common strategy^8^. This involves using statistical methods to estimate and fill in missing values, enabling subsequent data analysis. Imputation methods can be classified into naive imputation, feature-based imputation, and global-based imputation^7^. Deep learning model-based imputation methods have also been developed recently^9^.

However, most of the currently developed missing value imputation methods aim to construct a complete data matrix. This leads to the assumption that there should be a value for each missing place, without addressing the nature of the missing values. This overlooks the scenario that the missing values may truly reflect the absence of certain proteins in a sample. In fact, no/low expression in proteomics datasets may serve to distinguish disease activity^10^, including cancer progression^11^. Imputation can obscure the diagnostic value of such “hallmark” absence, which is often underappreciated and lacks comprehensive investigation.

Here, we first examined multiple proteomics datasets and substantiated that missing values (NAs) could segregate samples of different tissues of origin. To excavate the biological significance of NAs, we developed BIND (**B**iologically **I**nformative **N**A **D**econvolution), a machine learning workflow that deconvolves “biological” NAs indicating low/no expression and “technical” NAs resulting from experimental errors. We compiled BIND to a web server as a public tool that delineates missing values in proteomics, facilitating deep data exploration with interactive visualization. In both bulk and single-cell proteomics datasets, BIND identified the “hallmark absence” of unique proteins in cell lines and cancer extracellular vesicles (EVs). This helped to identify markers of hematologic cancer and cancer EV. Moreover, BIND-deconvolved NAs also improved the identification of protein complexes. Altogether, BIND represents a systematic tool that effectively delineates the nature of missing values and extracts biological insights.

## Materials and methods

### Data collection and acquisition

Four large-scale proteomics datasets were used in this study. Three of them are human cancer cell line datasets, CCLE949, NCI60, and CRC65. In addition, a human tissue extracellular vesicle dataset, HumanEV, was used to analyze the exosomes of cancer and normal tissues.

#### CCLE949

Quantitative DIA analysis was performed on 949 cell lines from 40 cancer types in 28 tissues^12^. The resulting dataset was derived from 6,864 DIA-MS runs acquired over 10,000 MS h, including HEK293T for all data acquisition cycles and quality control instrumentation. SWATH-MS measurements were performed on a 6600 TripleTOF system. A total of 6,692 human proteins were quantified.

#### NCI60 and CRC65

NCI-60 consists of 59 individual cancer cell lines from 9 different tissues (brain, blood and bone marrow, breast, colon, kidney, lung, ovary, prostate, and skin)^13^. These cells were lysed and digested with trypsin into a nanoLC-Orbitrap Velos mass spectrometer for analysis. Data-dependent acquisition (DDA) was used in positive ion mode. The CRC65 data contains proteomics datasets of 65 colorectal cancer cell lines with the same data acquisition process as NCI60^14^.

#### HumanEV

Extracellular vesicle (EV) protein samples from cancerous and normal tissues of various cancer patients^11^. Tissues were cut into small pieces and cultured for 24 hours in serum-free RPMI supplemented with penicillin and streptomycin. Conditioned media were processed for EV protein isolation and EV proteins were purified by sequential ultracentrifugation. Enriched EV protein samples were dried by vacuum centrifugation and re-dissolved for reduction, alkylation and trypsin digestion. Peptides were separated on a C18 column and the mass spectrometer (Q-Exactive, Q Exactive Plus, Q-Exactive-HF or Fusion Lumos) was operated in data-dependent acquisition (DDA) positive ion mode.

#### Single-cell proteomics (SCP)

Single-cell proteomics can provide proteomics data on a larger scale and with higher resolution^15^. We used a Hela and Jurkat cell line SCP dataset^16^. Hela and Jurkat cell cultures were performed in vitro, and single-cell fractionators were used to generate single cells. Cell lysis, protein denaturation, and digestion were performed using a mixture of trypsin and LysC digests. NanoLC separations were performed using an UltiMate 3000 RSLCnano system with a SPE column. MS analysis was performed using an Orbitrap Exploris 480 equipped with a Nanospray Flex Ion Source.

### Statistical analysis of missing values

Given a protein expression matrix *X*, where *x_i,u_* is the expression of protein *u* in sample *i*, we define the proportion of NA for a protein *u* as:

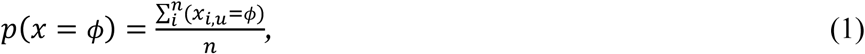

where we use *ϕ* to denote the missing value, the NA, in the protein matrix, and *n* is the sample number.

An NA proportion of 0 (*p*(*x*=*ϕ*)=0) indicates that the protein is present in all samples, while an NA proportion of 1 (*p*(*x*=*ϕ*)=1) means it is absent in all samples. This method allows for calculating NA proportions across the entire dataset or within specific groups, enabling a detailed analysis of missing data patterns for each protein.

We binarize the dataset by replacing NA with 0 and other values with 1, allowing for the treatment of the data as asymmetric binary data^17^, i.e., transfer *x_i,u_* to 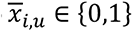. This enables the calculation of the Jaccard distance between samples, facilitating the analysis of similarity and dissimilarity in the presence of missing values^18^:

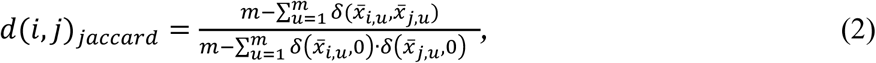

where *i* and *j* are two samples and *u* is proteins, *m* is the total number of proteins. *δ* is the Kronecker delta function and is defined as:

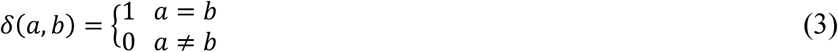

These calculated distances are then used for hierarchical clustering of the samples and the subsequent classification of missing values.

### Classification of missing values

We calculate the distance between two samples, either the Jaccard distance or the Euclidean distance:

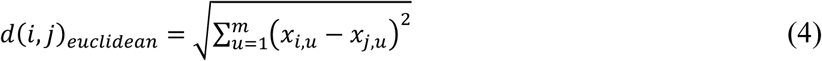

The gaussian kernel function is then applied to convert the distance *d*(*i,j*) between sample *i* and *j* to similarity:

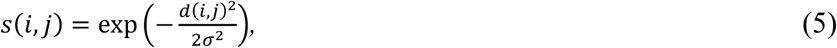

where *σ* is the function’s parameter to control the attenuation slowdown of similarity^19^.

We denote *y_i,u_* as the weighted expression value of the protein *u* in the *k* nearest samples of the sample *i*:

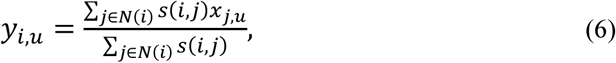

where *N(i)* is the *k* nearest neighborhood sample found by the sample distance. *x_j,u_* was set to 0 if it is originally a missing value *x_j,u_*=*ϕ*, ensuring that missing cases in neighbors are also considered.

Finally, we assign a label *l_i,u_* of “biological” (*ϕ^b^*) or “technical” (*ϕ^t^*) to each weighted expression *y_i,u_* with a manual threshold *γ*:

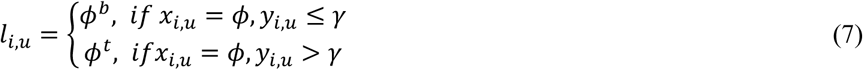

Next, we evaluate the performance of NA classification using simulated data from a complete dataset. For a large-scale dataset, all data with no missing values are used as a baseline. *a*% of the data is randomly removed from the smallest 1.2∗ *a*% of the data to simulate “biological” NAs. Then, *b*% of the remaining data is randomly removed to simulate “technical” NAs. The performance is assessed based on the accuracy of NA classification.

### Missing value-assisted protein-protein interactions analysis

Based on the analysis of specific individual proteins and to achieve a more sensitive analysis of protein interactions, we take into account the pattern of missing values in the two proteins in the calculation of the correlation coefficients. The Spearman correlation coefficients (SCC) of the two proteins *u* and *v* in the dataset were first calculated.

In BIND, we introduce a weighted correlation *ρ* based on *ρ_scc_* with a reward-penalty weighting scheme:

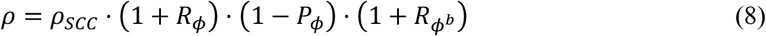

The *R_ϕ_* term rewards co-occurring *ϕ*s, *P_ϕ_* penalizes mismatched patterns, and *R*_bio_ provides an additional reward for ”biological NA” *ϕ^b^*s based on Equation (7):

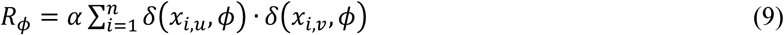

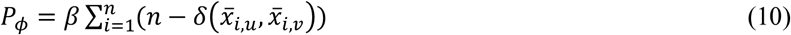

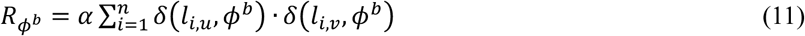

Here, *α* and *β* are user-defined reward and penalty coefficients, respectively, with default values of *α*=*β*=0.01. Finally, the *ρ* is adjusted to *ρ_BIND_* in the range -1 to 1:

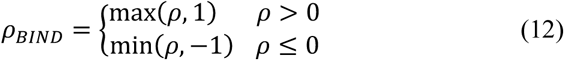

With *ρ_BIND_*, a more sensitive analysis of protein interactions can be realized. We demonstrate this from several perspectives in the results section.

### Clinical sample collection and processing

The research protocol received ethical approval from the Human and Artefacts Ethics Committee at City University of Hong Kong (approval number: HU-STA-00000410). All procedures involving clinical sample collection and experimental protocols adhered strictly to the principles outlined in the Declaration of Helsinki.

Whole blood specimens were obtained from clinical patients, with ages spanning from 10 to 82 years. The blood was transferred into anticoagulant-containing tubes with secure lids for centrifuging. EV was isolated by centrifugation, and its concentration was determined by measuring the absorbance at 280 nm using a spectrophotometer.

Samples were first reduced by TCEP and then alkylated by IAM (iodoacetamide). Proteins were digested with Trypsin/Lys-C Protease Mix. The digestion was acidified by 2% FA and then desalted by homemade C18 tips.

### LC-MS proteomic analysis

Dried samples were reconstituted in formic acid (FA) in water and loaded onto disposable EvoSep C18 EvoTips following the manufacturer’s instructions. Peptide elution was carried out at a high flow rate. Chromatographic separation utilized a predefined gradient method (44-minute runtime).

Protein samples were processed and injected into an Orbitrap Explori 480 mass spectrometer. Full MS scans were performed in data-dependent acquisition (DDA) mode. The raw mass spectrometry data were analyzed using Thermo Scientific Proteome Discoverer 2.4 (PD 2.4). MS/MS spectra were searched against the *Homo sapiens* UniProt database supplemented with common contaminants. Label-free quantification (LFQ) was conducted using the normalized abundance values generated by PD 2.4.

### Bioinformatics tool and web server implementation

The algorithm and function of BIND were compiled in R (version 4.3.2). The web tool was implemented based on JavaScript. Users can access and process their data freely in BIND without any login requirement or cookies through some popular web browsers. Users can choose to operate this tool on their computers by using our provided command-line scripts. The detailed user manual can be found in our GitHub repository (see Data availability).

## Result

### Characterization of missing values in cell line datasets

Large-scale proteomics studies typically generate many missing values (NAs). We examined NAs in three large-scale cancer cell line proteomics datasets: CCLE949, NCI60, and CRC65 (Supplementary Table S1). The CCLE949 dataset contains SWATH-based proteomics analysis of 949 common cancer cell lines. NCI60 had DDA proteomics data of 60 NCI cell lines, 20 of which are also analyzed in CCLE949. CRC65 contains 65 colorectal cancer cell lines by DDA proteomic analysis, providing the most targeted sample source. A common feature of studies is the lack of replicates, making it challenging to identify technical NAs.

We exploited binary heatmaps to visualize the global pattern of NAs (Figure 1A). Overall, CCLE949 contains 6692 proteins, and 487 proteins (7.3%) were detected in all 949 samples (no NA). In addition, 1808 (27%), 2337 (34.9%) and 2060 (30.8%) proteins were recorded as NA in < 20%, 20- 80% and > 80% of samples, respectively (Figure 1B, top panel). In comparison, NCI60 and CRC65 recorded more proteins (> 10,000) and showed substantially higher proportions of proteins without NAs (37.8% and 41.9%, respectively, Figure 1B, middle and down panel). This difference could be due to the sample numbers, heterogeneity, and MS performance.

**Figure 1.**
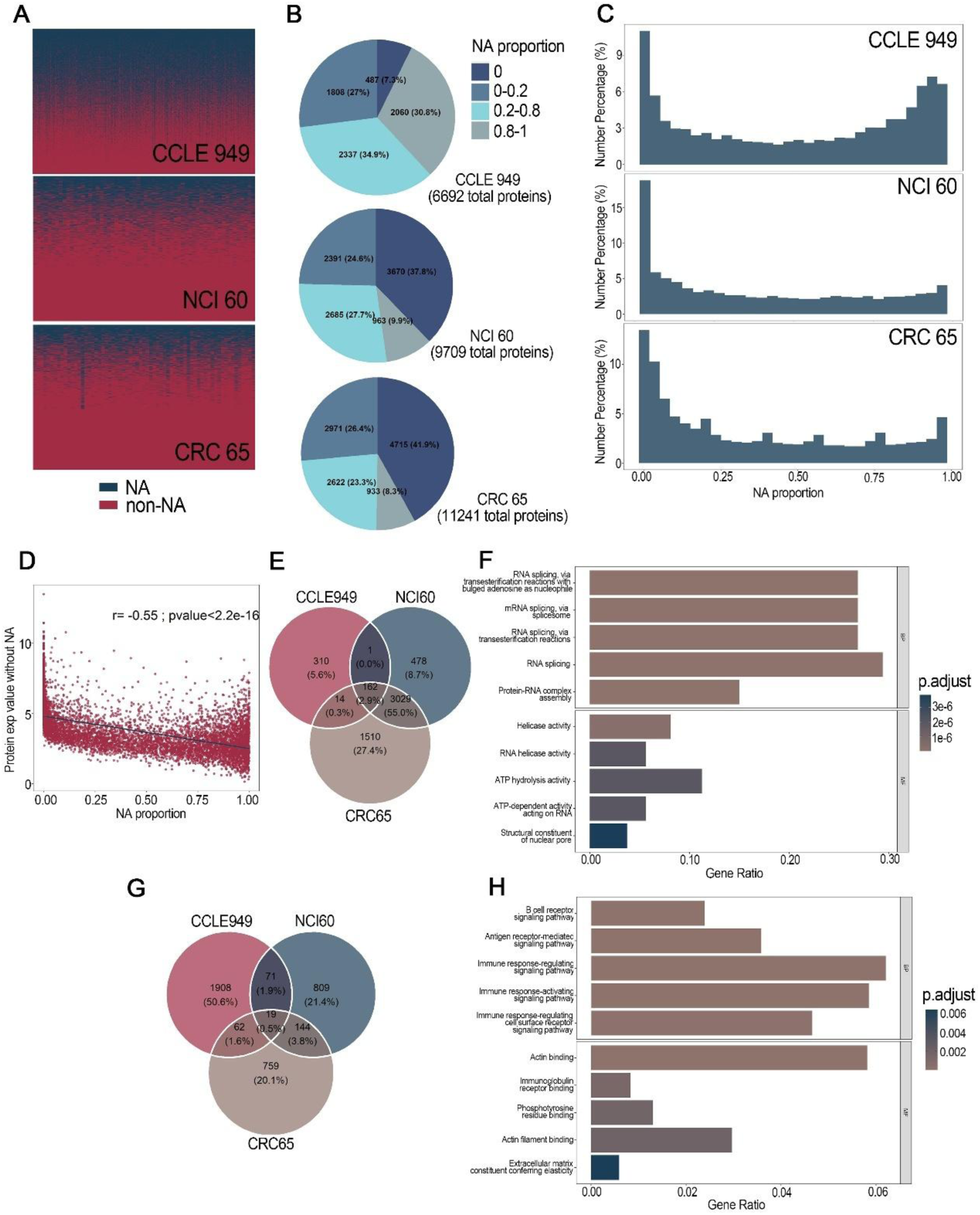
Characterization of missing values in three large-scale cell line datasets. (A) Binarized heatmap of missing values for human cancer cell line datasets CCLE949, NCI60, and CRC65. Blue: NA, Red: non-NA(expression value). (B) NA distribution pie plot of three cell line datasets. (C) NA proportion distribution histogram of the datasets. (D) Scatter plot of mean expression value to NA proportion of proteins in the CCLE949 dataset. (E) Venn diagram of never-missed proteins in three datasets, 162 proteins never missed in all 3 datasets. (F) GO enrichment analysis of never-missed proteins. (G) NA proportion more than 0.8 proteins in three datasets. (H) GO enrichment analysis of unique proteins with NA proportion >0.8 in the CCLE949 dataset.

To measure the magnitude of missingness, we utilized the “NA proportion” to represent the frequency at which a particular protein was recorded as NA in the dataset. Histograms indicated that all three datasets have more proteins with high and low NA proportions than in the middle range (Figure 1C). Notably, the protein NA proportions displayed a significant negative correlation with the mean expression level (*r = -0.55, p<2.*2e-16) (Figure 1D, Supplementary Figure S1). This is also observed among individual samples in which the high-abundance proteins tend to have low NA proportions (Supplementary Figure S2). This suggests that the frequency of missing values could reflect the expression level of the protein.

For the no-NA proteins, 162 were shared among all three datasets, suggesting their common essentialness (Figure 1E, Supplementary Table S5). Indeed, GO-term analysis showed that these proteins are enriched in housekeeping pathways, including spliceosome, helicase activity, and nucleocytoplasmic transport (Figure 1F). In contrast, the three datasets only shared 19 (0.5%) proteins with NA proportion > 80%, with CCLE949 having the most unique high-NA proteins (Figure 1G). GO-term analysis revealed that these proteins are enriched in immune- and cytoskeleton-related functions (Figure 1H). This is in line with the fact that CCLE949 included heterogeneous samples of hematopoietic and solid cancer cell lines of distinct culture conditions. These results indicate that BIND-mediated NA analyses could point to the biological significance of proteins, including unique expression patterns and functional status. For further investigations, we next focused on the CCLE949 dataset with the most heterogeneous tissue types.

### NA patterns discern cells from different tissue of origin

The cell lines in CCLE949 were derived from various tissues (Supplementary Figure S3). To focus on distinctive tissue origins, we investigated the NA patterns of the hematopoietic, lung, breast, large intestine, and stomach cancer cell lines. Clustering analysis based on the NA proportions revealed that the solid cancers are highly correlated in their NA patterns but are distinct from the hematopoietic cancers (Figure 2A). In lung and hematopoietic cancers with the largest sample size, we also observed more accumulation of high- or low-NA proteins than proteins with middle-level NA proportions (Figure 2B). Notably, hematopoietic cancers have more proteins with an NA proportion > 0.9, indicating greater proteome variations than the lung cancer cells.

**Figure 2.**
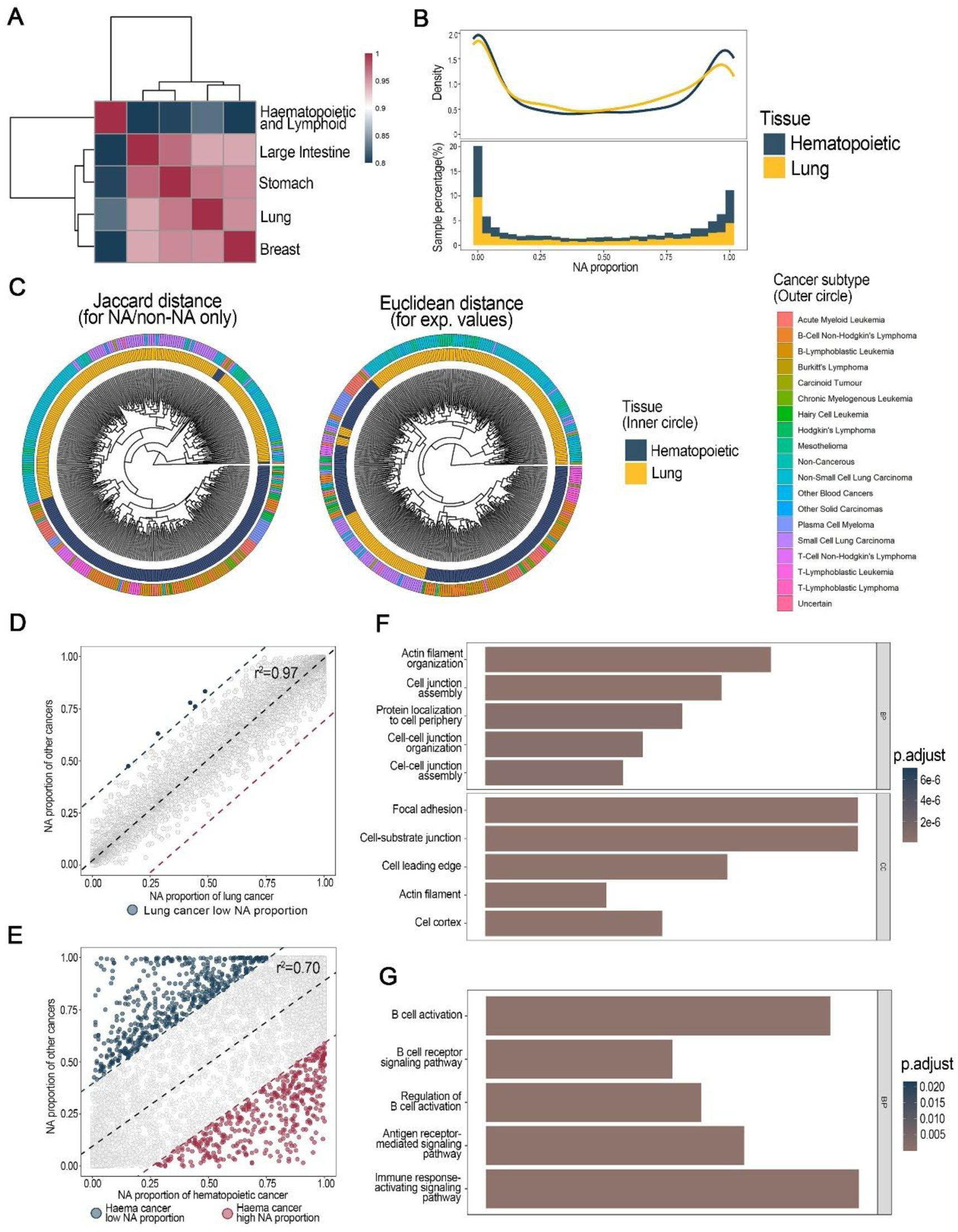
Cell lines from different tissue origins showed different patterns of missing values. (A) Correlation heatmap of NA proportions in hematopoietic, large intestine, stomach, lung, and breast tissue. (B) NA proportion distribution density (up) and histogram(down) of hematopoietic and lung tissues. (C) Sample clustering using Jaccard distance (left) and Euclidean distance (right). Binarized Jaccard distance achieved better separation of origin tissues. (D) Scatter plot of NA proportions in other tissues to NA proportions in lung cancer. Lung cancer had a similar NA pattern to other tissues. (E) Scatter plot of NA proportions in other tissues to NA proportions in hematopoietic tissue. Hematopoietic cancer showed significant differences from other tissues. Red point: proteins with high NA proportions in hematopoietic cancer, but low NA proportions in other tissues; Blue point: proteins with low NA proportions in hematopoietic cancer, but high NA proportions in other tissues. (F) GO enrichment of highly missing proteins in hematopoietic cancer. (G) GO enrichment of seldom missing proteins in hematopoietic cancer.

To investigate whether NA could serve as a specific feature that classifies tissue types/subtypes, we binarized the dataset by assigning “0” to NAs and “1” to the detected values (Supplementary Table S6). We performed clustering analysis of the hematopoietic and lung cancer samples with Jaccard distance, which is tailored for binary data. The results showed excellent separation of the two cancer types with subtype stratification of the lung cancer cell lines (Figure 2C, left panel). Interestingly, NA-based binary clustering produced a better outcome than clustering with Euclidean distance after removing all NAs (Figure 2C, right panel). This result illustrates that NAs could inform specific absence of proteins and differentiate tissue types.

To further investigate the NA features of the two cancer types, we compared the NA proportion of their proteomes with other cancer types (Figure 2D-E). Scatter plots revealed a good correlation between the NA proportions of the lung cancer proteome and the other types of solid tumor (Figure 2D). In contrast, the hematopoietic cancers displayed a poor correlation in protein NA proportions with the other cancer types (Figure 2E). We then analyzed the proteins with a substantial difference (exceeding +/- 30% NA proportions) between the hematopoietic cancers and the solid-tumor cancer types (blue and red dots in Figure 2E). GO enrichment analysis showed that proteins with high NA proportions in hematopoietic cancers were enriched in focal adhesion and cell junction assembly (Figure 2F). This is in line with the absence of such structures in hematopoietic cell lines, which are typically in suspension cultures. In contrast, proteins with low NA proportions in hematopoietic cancers were mainly involved in immune cell activation and immune response signaling, consistent with their absence from the solid-tumor cell lines (Figure 2G). Our results indicate that the NA patterns of the cell line proteomes could discern the tissue-specific presence and absence of proteins.

### BIND deconvolves NAs and identifies the unique absence of proteins in human cancer cells

In addition to representing low/no expression, NAs can also be due to experimental errors. To excavate the biological information carried by the missing values, it is important to deconvolve the nature of NAs, which we classify here into two categories: “biological” NAs that reflect no/low protein expression, and “technical” NAs introduced by non-biological errors. We developed the **B**iologically **I**nformative **N**A **D**econvolution (BIND) algorithm that applies an adaptive k-nearest neighbor-based algorithm for determining the nature of NAs (Figure 3A, see Methods for details). Briefly, BIND dynamically adjusts the Gaussian kernel bandwidth of each sample according to the local neighborhood structure to compute context-aware similarity weights. These weights are then integrated with simultaneously captured expression values and co-occurrences of missing values within the neighbors. Finally, BIND determines “biological” or “technical” NAs that are below or above an optimized threshold γ, respectively.

**Figure 3.**
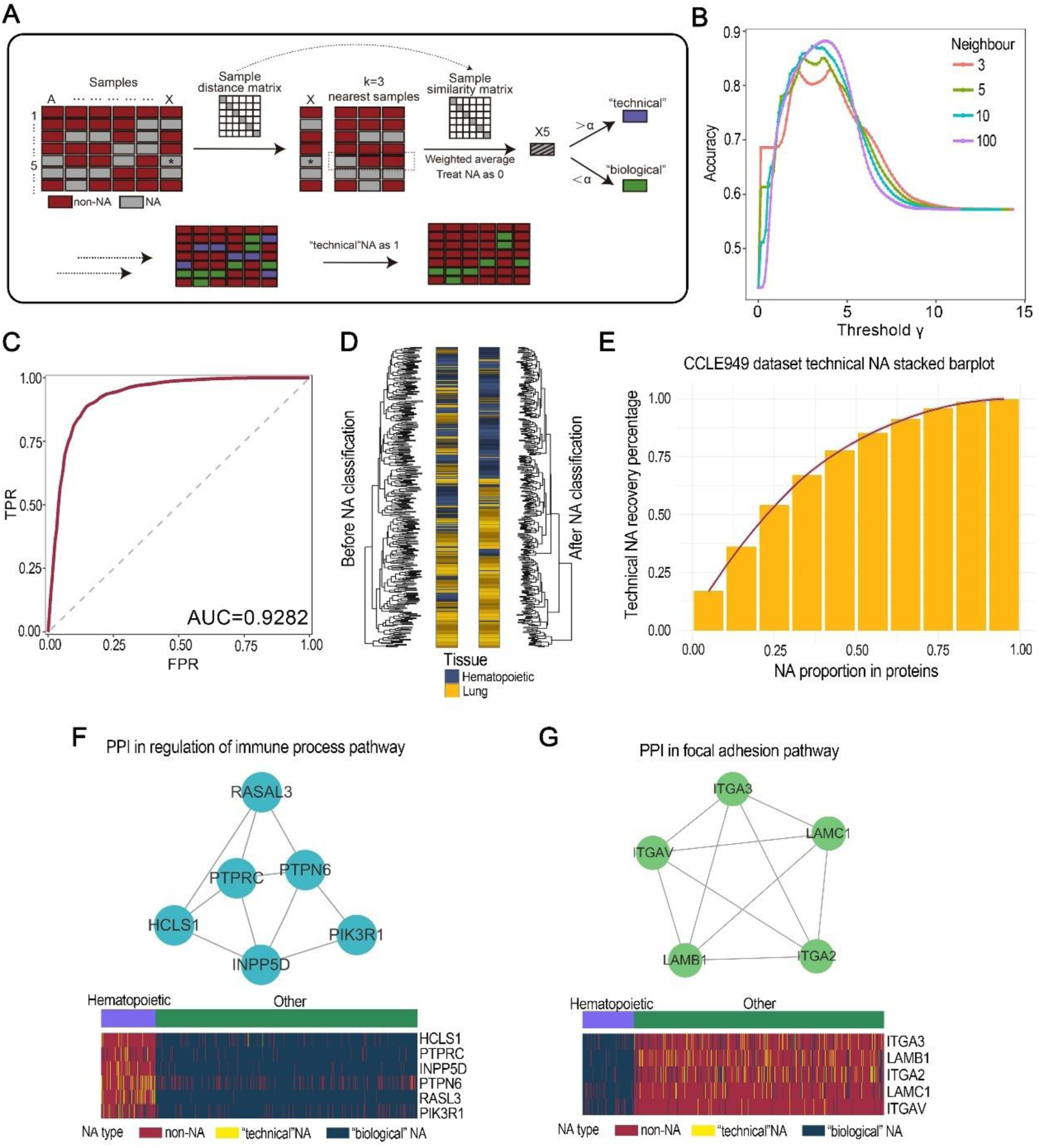
NA source deconvolution uncovered unique proteins from human cancer cell line data. (A) Workflow of NA classification. The figure illustrates the classification process of trying to determine the category of missing values at position X5. (B) Variation of NA classification accuracy with different parameters in the simulation dataset, we chose k=10 and threshold =3 for further study in the CCLE949 dataset. (C) When k=10, the AUC of NA classification achieved at 0.9282. (D) Binarized clustering of the simulation dataset before and after NA classification. The “technical” NAs were designated as 1, as an expression value should have existed there. (E) A stacked histogram of “technical” NA, with overall NA proportion in the horizontal coordinate. (F) BIND helps to recover proteins that are exclusive to hematopoietic cancer. This protein complex is involved in the immune process pathway. (G) BIND helps restore proteins that are unique to other tissues. This protein complex is involved in focal adhesion pathways.

We exploited the CCLE949 dataset and generated a collection of simulated “biological” and “technical” NAs to optimize BIND. Briefly, a fraction of low-expression proteins was removed to create “biological” NAs, and an additional fraction of quantified proteins was discarded to mimic “technical” NAs. Reiterative modeling demonstrated that the performance of BIND improves with increasing k (number of neighboring samples) and varying threshold γ (Figure 3B). For the simulated dataset, BIND determined the optimal parameters (k = 10 and γ = 3) with an AUC = 0.9282 in deconvolving the two types of NAs (Figure 3C, Supplementary Figure S4). Notably, the NA deconvolution substantially improved distinguishing hematopoietic and lung cancer samples by binary transformation of the dataset (Figure 3D). This suggested that BIND could ameliorate the interference of the “technical” NAs during multivariate analysis of proteomic data.

We next applied BIND to the CCLE949 dataset and evaluated the performance. We observed a fast increase in the technical NA identification as the NA proportion of proteins increases (Figure 3E). For proteins with 50% NA proportion, 80% of the technical NAs were recovered. Notably, hematopoietic cancer and other cancer types have collections of proteins with distinct biological-to-technical NA ratios, revealing more proteins of unique absence/presence (Supplementary Figure S5). For example, a large number of NAs of HCLS1, PTPRC, INPP5D, PTPN6, RASL3, and PIK3R1 are determined as “technical” ones in hematopoietic cancer samples, but not in other cancer types (Figure 3F). This suggests the unique presence of these proteins in hematopoietic cells, in line with their immune-regulation functions. In contrast, ITGA2, ITGA3, ITGAV, LAMB1 and LAMC1 have most NAs determined as “biological” ones in hematopoietic samples and as “technical” ones in other cancer types (Figure 3G). This aligns well with their functional implications in the focal adhesion pathway, which should be insignificant in hematopoietic samples. Interestingly, these proteins have intricate PPIs, which prompted us to investigate whether BIND could help improve PPI analyses.

### BIND enables more sensitive co-expression analysis in identifing protein-protein interactions

Protein expression covariance has been exploited for in protein-protein interaction (PPI) analysis. However, the presence of NAs has been a significant barrier for such analysis. We propose that co-appearance as biological NAs is a rewarding factor when calculating expression covariance of two proteins. Therefore, following deconvolving biological NAs, BIND implements a missing value-weighted approach to generate *ρ_BIND_*, an NA-assisted correlation coefficient to improve PPI analysis (Figure 4A, see methods for details).

**Figure 4.**
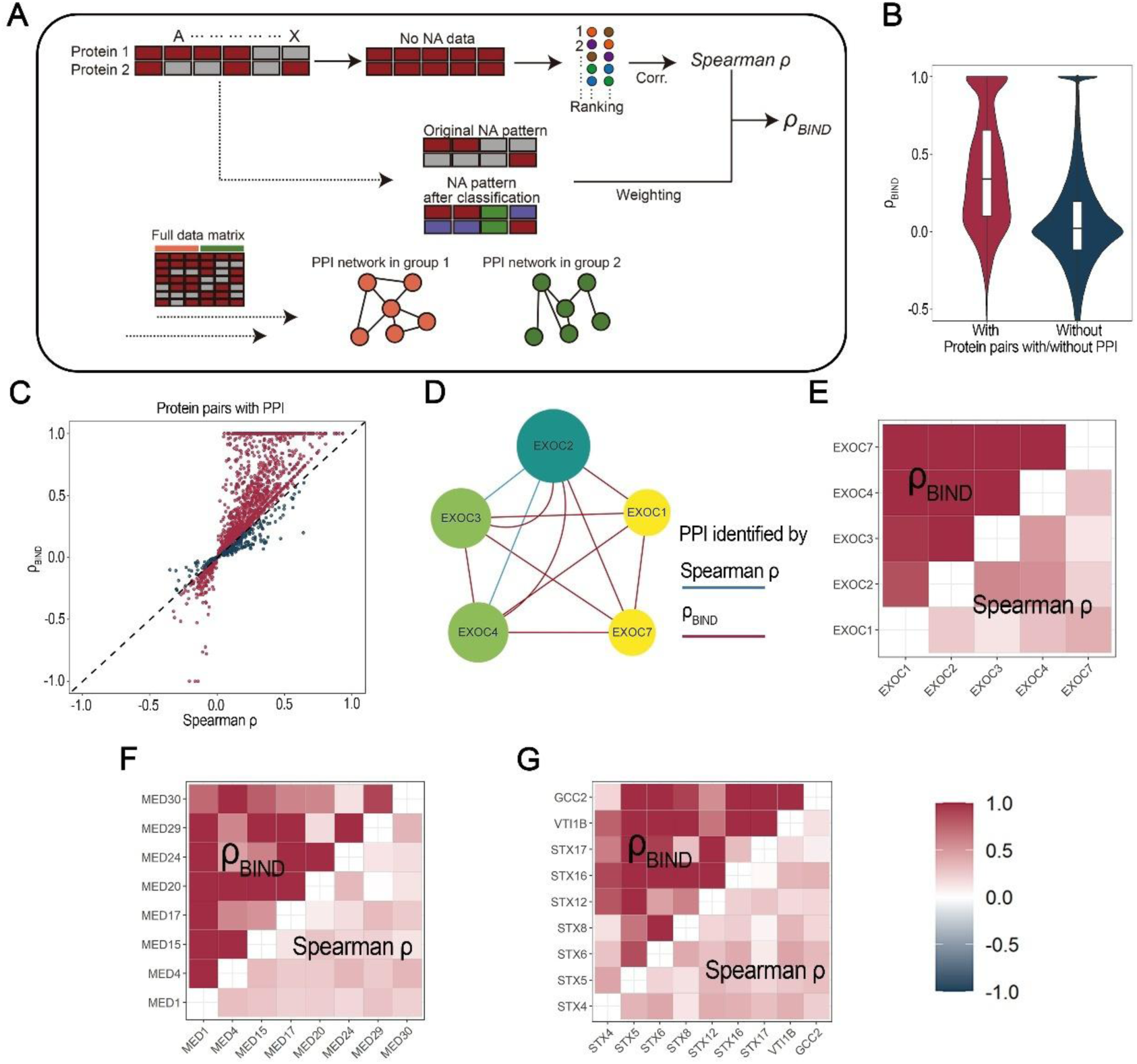
NA-assisted protein co-expression analysis identified protein-protein interactions more sensitively. (A) Workflow of NA-assisted protein co-expression analysis. This approach takes missing value patterns into account in Spearman correlation analyses, rather than just using non-missing data. (B) Distribution and mean value of *ρ_BIND_* in protein pairs with and without PPIs. (C) Scatter plot of protein pairs with PPI, *ρ_BIND_* to Spearman *ρ*. Red: |*ρ_BIND_|* larger than |Spearman *ρ|*; Blue: |*ρ_BIND_|* smaller than |Spearman *ρ|*. (D) PPI in EXOC protein complex identified by Spearman *ρ* and *ρ_BIND_*, 0.5 as threshold. *ρ_BIND_* identified all 10 interactions. (E) Heatmap of Spearman *ρ* and *ρBIND* in EXOC complex PPI identification. (F-G) Heatmap of Spearman *ρ* and *ρ_BIND_* in MED and STX complexes. *ρ_BIND_* showed more sensitive identification.

We first analyzed the hematopoietic cancer samples in the CCLE949 dataset. Human protein interactions were obtained from the STRING database, and 3556 pairs of physical interactions with PPI scores greater than 0.9 were considered “with PPI”. The mean value of *ρ_BIND_* for protein pairs with PPI was higher than that for protein pairs without PPI (Figure 4B). For 79.1% of these PPI pairs, *ρ_BIND_* yielded high association coefficient values than the conventional Spearman *ρ* (Figure 4C, Supplementary Figure S6). Moreover, with > 0.5 as the threshold, *ρ_BIND_* values established 10(100%) interactions among the EXOC complex, whole conventional Spearman *ρ* only detected 2(20%) interactions (Figure 4D-E). This is also the case for MED and STX (Fgiure 4F-G), and others. In addition, for the known protein complexes in the CORUM database, the ρBIND within complexes is higher than between complexes (Supplementary Figure S7). This further demonstrates the ability of *ρ_BIND_* to characterize tight protein-protein associations. In summary, *ρ_BIND_* could further extarct information from missing values, and improve the PPI analysis with expression covariation.

### BIND identified protein markers of extracellular vesicles in human cancers

We applied BIND to another large-scale proteomic dataset, an EV dataset from human cancers and normal tissues. It contains a high number of missing values and helps to evaluate the performance of BIND in this scenario. Generally, the EV dataset is more irregular than the cell line dataset (Figure 5A), containing more proteins with an NA proportion close to 1 (Figure 5B). After applying BIND to identify biological NAs and NA-guided binary transformation (Figure 5C), the resulted data was effective in clustering analysis to discriminate between cancer and normal EV samples (Figure 5D).

**Figure 5.**
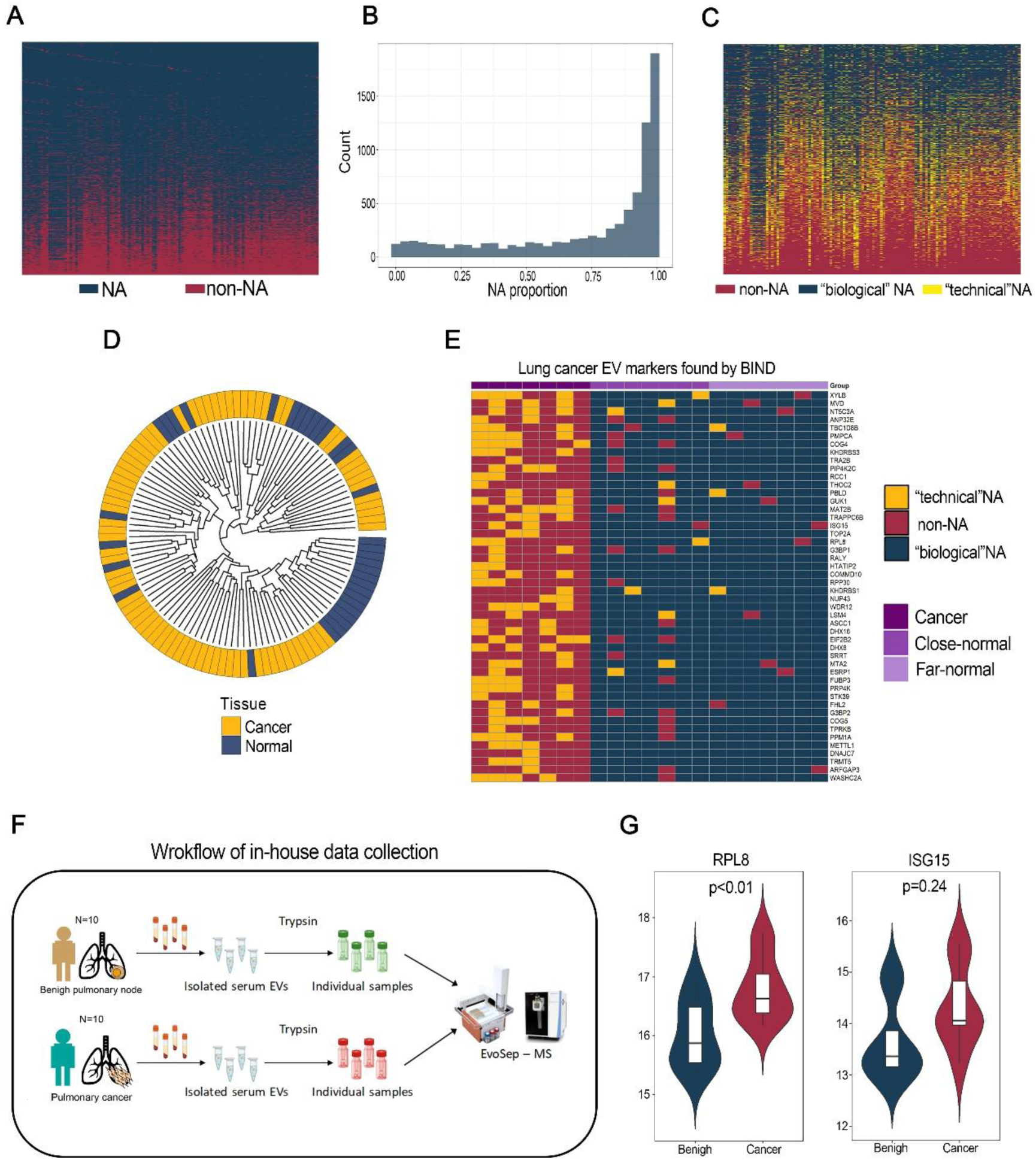
BIND identified protein markers of extracellular vesicles in human cancers. (A) Missing value heatmap of human cancer/normal EV dataset. (B) NA distribution of EV dataset. (C) NA heatmap after classification. Red: expression value; Blue: “biological” NA; Yellow: “Technical” NA. (D) Clustering of Binarized EV dataset after NA classification. (E) NA classification helps to identify lung cancer EV protein markers. (F) Workflow of in-house proteomic data collection. We obtained serum EV proteomes from 10 patients with lung cancer and 10 patients with benign lung nodules. (G) Expression values of RPL8 and ISG15 protein in the in-house EV data. Statistical significance was obtained by Student’s t-test.

We next analyzed EV proteins from three paired sample groups in the dataset: lung cancer tissues, close normal tissues, and far normal tissues, and identified multiple newly discovered lung cancer EV proteins (Figure 5E).

In order to validate the lung cancer EV markers obtained by BIND, we performed in-house data collection. We performed proteomics analysis on serum EVs collected from 10 patients diagnosed with lung cancer and 10 patients with benign lung nodules (Figure 5F). Among the lung cancer EV proteins discovered by BIND, higher expressions of RPL8 and ISG15 were observed in malignant EVs (Figure 5G). These results showed that BIND could increase the sensitivity to discover potential biomarkers by delineating the nature NAs.

### A user-friendly BIND web server

Next, we compiled BIND into a web-based tool for researchers of different computational skill levels to explore and process NAs in large proteomics datasets interactively. It executes four functions: (1) Data input. A proteomics dataset containing missing values is imported with the grouping information. (2) Characterization of missing value patterns in the dataset. This part analyzes the pattern of missing values in the dataset and the differences of missing values in different subgroups through various visualization methods, such as heatmaps, pie charts, etc. BIND analyzes the NA frequency of proteins and performs sample clustering based on the binary transformation of missing values. (3) Classification of NAs. BIND differentiates “biological” from “technical” NAs by integrating features of neighboring samples (see Methods for details). This deconvolves the NA natures and helps interpret the biological significance. (4) NA-weighted protein interaction analysis. BIND generates weighting factors based on biological NAs to improve the abundance covariance analysis for more sensitive and precise mapping of protein complexes.

The BIND website provides pages including home page, user tutorial, example data view, task submission, task query, and result view (Figure 6A). In a typical workflow, users first enter the Submit Task page, where they upload data (including expression matrices, grouping information, etc.) and define the parameters of the task. A table file of example data is also available for download. Next, after the completion of a task, users can access the detailed results page, which contains readable and interactive graphs. The web page works on various popular computer systems (Windows, Mac, Linux) and popular web browsers (Chrome, Edge, Firefox, Safari, etc.) (Supplementary Table S9).

**Figure 6.**
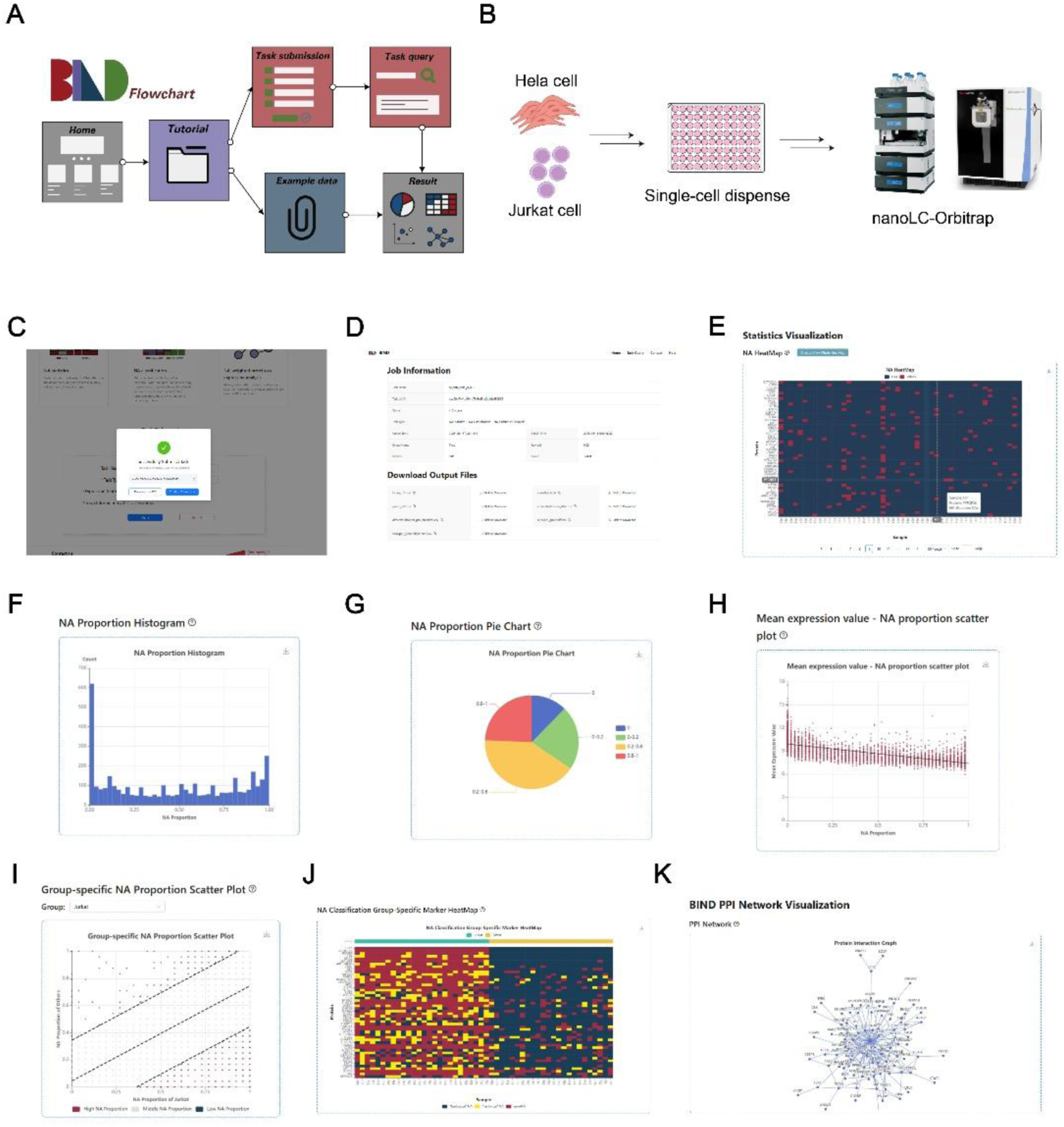
The online web server for BIND. (A) Flowchart of the BIND website. (B)Single-cell proteomics process of Hela and Jurkat cells. (C) Task submission page. (D)Task information page. After task completion, the user can view the result on an interactive page. (E)Binary heatmap showing NA and expression values. (F) Histogram of the distribution of NA proportion. (G) NA proportion’s pie chart. (H) Scatterplot of NA proportion vs. expression of this protein. The expression of the protein shows a negative correlation with the proportion of its missing. (I) Group comparison scatter plot of NA proportion (Hela vs Jurkat). The red dots in the plot are the proteins present in high proportion in Jurkat cells. (J) The Jurkat cell protein marker heatmap discovered by BIND. (K) Unique PPIs for Jurkat cells in the regulation of immune response pathway identified by BIND.

### Analyzing single-cell proteomics data on the BIND web server

Single-cell proteomics (SCP) enables the analysis of protein expression at the individual cell level, providing insights into cellular heterogeneity that cannot be captured by bulk-scale studies. SCP has developed rapidly recently for understanding complex biological mechanisms^15^. We expect that deconvolution of NAs in the SCP datasets could provide additional insights. Therefore, we examined the operation of the BIND web server utilizing an SCP dataset of two cell lines^16^ (Figure 6B). After tusk submission, a UUID is generated for task query (Figure 6C). Upon completion, visualization/download of results can be performed on the task information page (Figure 6D). The results include a heatmap of NA distribution (Figure 6E) as well as histograms and pie charts of NA proportions (Figure 6F-G). Similar to the analysis of bulk samples, we observed a negative correlation between NA proportion and expression of a protein (Figure 6H). Selective analysis could reveal proteins that are uniquely missing/present in Jurkat cells relative to Hela cells (Figure 6I). Following the NA deconvolution of SCP data, BIND identified protein markers of Jukart cells (Figure 6J), including LCP1 (Lymphocyte cytosolic protein 1) and EVL (Enah/Vasp-like). These protein markers are implicated in pathways such as cell leading edge and phagocytic vesicle (Supplementary Figure S8). Moreover, BIND improved PPI analysis and identified a GDI1 (GDP dissociation inhibitor 1)-centred complex that is unique to Jurkat cells (Figure 6K).

Overall, the BIND server represents a programming-free and interactive web-based tool for NA deconvolution. With the growing amount of large-scale proteomics and SCP data, it is expected to further our understanding of complex biological processes and disease mechanisms.

## Discussion

As with much large-scale biological data, the problem of missing values in proteomics data is a headache. Deeper protein identification brings more, not fewer, missing values as technology advances. A variety of imputation methods for proteomic data have been developed to fill into a complete data matrix for data analysis. However, these methods usually assume that the data matrix should be complete and needs to fill it in. Missing data are common and biologically logical, and simply filling them in may obscure the underlying biological value. BIND provides a novel analytical framework for viewing proteomics data from the perspective of the missing values, aiming to excavate the biological significance from the patterns of missing values.

### Algorithm and framework design for BIND

Biological samples from different sources differ in the presence and absence of proteins. However, few systematic comparisons have been made of differences in the proportion of missing proteins between different groups in large-scale proteomics datasets. An early study identified unique proteins in patients with rheumatoid arthritis by proportion of missing, but this approach has not been studied in greater depth^10^. Our results showed that there were significant differences in the proportion of missing values in cancer cell lines of different tissue origins by abandoning the consideration of expression value information, and that potential biological information could be mined by this process. This prompts us to consider the biological significance of missing values further. Missing values were categorized into three types, Missing completely at random (MCAR), Missing at random (MAR), and Missing not at random (MNAR)^20^. Although the missing value types are well recognized, few consider these scenarios during data processing. This may be due to the fact that we cannot exactly know the origin behind the missing values in a dataset. A recent study explored a method for differentiating missing value types based on the distribution of protein expression, but the method has only one uniform type for each protein, which remains deficient^21^. Instead of specifying whether a missing value is MNAR or MAR, BIND summarizes missing values into two scenarios, “biological” and “technical”, and is able to obtain the type of missing value for each missing position. Our method is based on the screening of samples that are neighbors of the sample in which the protein is located. This approach is effective and highly interpretable. One of the important difference of our proposed NA classification method compared to the common knn missing value imputation method was that the missing values in the dataset were treated as 0 during the computation, instead of ignoring them^22^. This is possible because our goal is to classify them, not to recover the expression value there, which ensures that the missingness of the neighbors is taken into account. Since “technical” NA, which we defined as due to a random technical problem, implied that there should have been a value there, our approach was not to impute the expression but simply to consider it as a non-missing value to be taken into account in the analysis of the missing value pattern. The results showed that this approach was effective in discovering, for instance, protein markers, and could be useful in different biological scenarios. At the protein-protein interaction level, we took the pattern of missing values into account in the Spearman correlation analysis, which is calculated by ranking data and has better robustness. The use of the PPI database to recover missing proteins in the dataset has been reported^23^. The STRING database provides known information on protein-protein interactions, which helps us to assess the reliability of our results.

### Biological significance found by BIND

By extracting the biological value of the missing values, BIND identified unique proteins and protein-protein interactions. We first revealed the biological functions of proteins that rarely showed missing based on NA patterns, which are involved in important pathways, such as RNA splicing, helicase activity, etc. This illustrated the significance of extracting biological insights from patterns of missing values and encouraged us to discover tissue-specific patterns of protein missingness. Further, after applying BIND on the cell line dataset, we found that BIND restores unique proteins in hematopoietic cancers that are present in the immune process pathway. Among them, HCLS1 is a hematopoietic cell-specific protein that is the substrate of the antigen receptor-coupled tyrosine kinase^24^. INPP5D plays a role in immune system homeostasis, neutrophil migration, etc^25^. PTPN6, although expressed in a wide range of cell lines, was significantly more expressed in hematological cancer cell lines than in other cell lines^26^. BIND recovered it in the majority of hematopoietic cancer cell lines, but little in other cell lines, although some of them expressed PTPN6. Similar patterns were observed in the PIK3R1 protein^27^. This suggests that BIND is sensitive to tissue type differences. On the other hand, LAMB1 and LAMC1 are subunits of laminin ^28,29^. They are utilized in the construction of extracellular matrix components and are minimally expressed in hematopoietic cancers. ITGA2, ITGA3, and ITGAV are receptors for fibronectin, laminin, etc., which are also widely present in various tissues other than hematopoietic cancers^30^. BIND also identified EXOC, MED, and STX complexes in hematologic cancers by PPI analysis. EXOC and STX play a role in vesicle transport, while MED plays a role in the regulation of RNA polymerase II^31–33^. In lung cancer EV, we validated the potential of RPL8 and ISG15 as markers, and expect subsequent research to further reveal their roles in lung cancer.

We provide biology researchers unfamiliar with programming a web server for data analysis. The web server has clean and clear pages and provides detailed instructions for operation. Its data analysis process is highly modular and highly interactive, allowing users to extract biological information of interest. We demonstrated the reliability of BIND for single-cell data analysis by testing a single-cell proteomics dataset on this web server. BIND could robustly analyze single-cell proteomics data, and identified characteristic protein markers as well as important protein-protein interactions in Jurkat cells.

In conclusion, BIND jumps out of the traditional missing value imputation process and provides unique insights into missing value-centered proteomics data analysis. We anticipate that BIND can facilitate in-depth analysis of proteomics data and contribute to the solution of the missing value problem. We also hope that the data analysis concept of “extracting value from the missing” will become more widely applied.

## Data Availability

All data and code related to this paper have been uploaded to GitHub repositories (https://github.com/guowh1999/BIND) and online databases. A user tutorial is provided. The address of the BIND online web server is https://bind.fengslab.com/. The public proteomics data used in the paper have been attached in Supplementary Tables to help researchers access them quickly.

## Supplementary Data

Supplementary tables and figures are provided.

## Funding

This work is supported by.

## Notes

### Competing Interest Statement

The authors have declared no competing interest.

## References

1 Zhang, Y. Y., Fonslow, B. R., Shan, B., Baek, M. C. & Yates, J. R. Protein Analysis by Shotgun/Bottom-up Proteomics. Chemical Reviews 113, 2343–2394 (2013). 10.1021/cr3003533

2 Li, M. B. & Smyth, G. K. Neither random nor censored: estimating intensity-dependent probabilities for missing values in label-free proteomics. Bioinformatics 39 (2023). 10.1093/bioinformatics/btad200

3 Vanderaa, C. & Gatto, L. Revisiting the Thorny Issue of Missing Values in Single-Cell Proteomics. Journal of Proteome Research 22, 2775–2784 (2023). 10.1021/acs.jproteome.3c00227

4 Pedersen, A. B. et al. Missing data and multiple imputation in clinical epidemiological research. Clinical Epidemiology 9, 157–165 (2017). 10.2147/clep.S129785

5 Lazar, C., Gatto, L., Ferro, M., Bruley, C. & Burger, T. Accounting for the Multiple Natures of Missing Values in Label-Free Quantitative Proteomics Data Sets to Compare Imputation Strategies. Journal of Proteome Research 15, 1116–1125 (2016). 10.1021/acs.jproteome.5b00981

6 Gardner, M. L. & Freitas, M. A. Multiple Imputation Approaches Applied to the Missing Value Problem in Bottom-Up Proteomics. International Journal of Molecular Sciences 22 (2021). 10.3390/ijms22179650

7 Kong, W. J., Hui, H. W. H., Peng, H. & Bin Goh, W. W. Dealing with missing values in proteomics data. Proteomics 22 (2022). 10.1002/pmic.202200092

8 Bramer, L. M., Irvahn, J., Piehowski, P. D., Rodland, K. D. & Webb-Robertson, B. J. M. A Review of Imputation Strategies for Isobaric Labeling-Based Shotgun Proteomics. Journal of Proteome Research 20, 1–13 (2021). 10.1021/acs.jproteome.0c00123

9 Webel, H. et al. Imputation of label-free quantitative mass spectrometry-based proteomics data using self-supervised deep learning. Nature Communications 15 (2024). 10.1038/s41467-024-48711-5

10 McGurk, K. A. et al. The use of missing values in proteomic data-independent acquisition mass spectrometry to enable disease activity discrimination. Bioinformatics 36, 2217–2223 (2020). 10.1093/bioinformatics/btz898

11 Hoshino, A. et al. Extracellular Vesicle and Particle Biomarkers Define Multiple Human Cancers. Cell 182, 1044-+ (2020). 10.1016/j.cell.2020.07.009

12 Goncalves, E. et al. Pan-cancer proteomic map of 949 human cell lines. Cancer Cell 40, 835-+ (2022). 10.1016/j.ccell.2022.06.010

13 Gholami, A. M. et al. Global Proteome Analysis of the NCI-60 Cell Line Panel. Cell Reports 4, 609–620 (2013). 10.1016/j.celrep.2013.07.018

14 Frejno, M. et al. Proteome activity landscapes of tumor cell lines determine drug responses. Nature Communications 11 (2020). 10.1038/s41467-020-17336-9

15 Tan, Y. C., Low, T. Y., Lee, P. Y. & Lim, L. C. Single-cell proteomics by mass spectrometry: Advances and implications in cancer research. Proteomics 24 (2024). 10.1002/pmic.202300210

16 Sanchez-Avila, X. et al. Easy and Accessible Workflow for Label-Free Single-Cell Proteomics. Journal of the American Society for Mass Spectrometry 34, 2374–2380 (2023). 10.1021/jasms.3c00240

17 Konecny, J. & Trnecka, M. Boolean matrix factorization for symmetric binary variables. Knowledge-Based Systems 279 (2023). 10.1016/j.knosys.2023.110944

18 Kryszkiewicz, M. in 16th Asian Conference on Intelligent Information and Database Systems (ACIIDS). 341–349 (2024).

19 Feng, X. K., Chen, L. X., Wang, Z. S. & Li, S. C. I-Impute: a self-consistent method to impute single cell RNA sequencing data. Bmc Genomics 21 (2020). 10.1186/s12864-020-07007-w

20 Liu, M. Y. & Dongre, A. Proper imputation of missing values in proteomics datasets for differential expression analysis. Briefings in Bioinformatics 22 (2021). 10.1093/bib/bbaa112

21 Lin, W. Q. et al. omicsMIC: a comprehensive benchmarking platform for robust comparison of imputation methods in mass spectrometry-based omics data. Nar Genomics and Bioinformatics 6 (2024). 10.1093/nargab/lqae071

22 Lalande, F. & Doya, K. in 15th International Conference on Similarity Search and Applications (SISAP). 3–10 (2022).

23 Kong, W. J. et al. PROTREC: A probability-based approach for recovering missing proteins based on biological networks. Journal of Proteomics 250 (2022). 10.1016/j.jprot.2021.104392

24 Skokowa, J. et al. Interactions among HCLS1, HAX1 and LEF-1 proteins are essential for G-CSF-triggered granulopoiesis. Nature Medicine 18, 1550–U1157 (2012). 10.1038/nm.2958

25 Metzner, A. et al. Reduced proliferation of CD34+ cells from patients with acute myeloid leukemia after gene transfer of INPP5D. Gene Therapy 16, 570–573 (2009). 10.1038/gt.2008.184

26 Tartey, S. et al. Ets-2 deletion in myeloid cells attenuates IL-1α- mediated inflammatory disease caused by a Ptpn6 point mutation. Cellular & Molecular Immunology 18, 1798–1808 (2021). 10.1038/s41423-020-0398-7

27 Nguyen, T. et al. Human PIK3R1 mutations disrupt lymphocyte differentiation to cause activated PI3Kδ syndrome 2. Journal of Experimental Medicine 220 (2023). 10.1084/jem.20221020

28 Lee, H. et al. Upregulation of LAMB1 via ERK/c-Jun Axis Promotes Gastric Cancer Growth and Motility. International Journal of Molecular Sciences 22 (2021). 10.3390/ijms22020626

29 Ye, G. X. et al. Lamc1 promotes the Warburg effect in hepatocellular carcinoma cells by regulating PKM2 expression through AKT pathway. Cancer Biology & Therapy 20, 711–719 (2019). 10.1080/15384047.2018.1564558

30 Nones, K. et al. Genome-wide DNA methylation patterns in pancreatic ductal adenocarcinoma reveal epigenetic deregulation of SLIT-ROBO, ITGA2 and MET signaling. International Journal of Cancer 135, 1110–1118 (2014). 10.1002/ijc.28765

31 Gromley, A. et al. Centriolin anchoring of exocyst and SNARE complexes at the midbody is required for secretory-vesicle-mediated abscission. Cell 123, 75–87 (2005). 10.1016/j.cell.2005.07.027

32 Hamasaki, M. et al. Autophagosomes form at ER-mitochondria contact sites. Nature 495, 389–393 (2013). 10.1038/nature11910

33 Zhang, Y. T., Qin, P. F., Tian, L. L., Yan, Y. G. & Zhou, Y. L. The role of mediator complex subunit 19 in human diseases. Experimental Biology and Medicine 246, 1681–1687 (2021). 10.1177/15353702211011701

