## Supplementary Figure S1-S8 for "Biologically Informative NA Deconvolution (BIND) excavates hidden features of the proteome from missing values in large-scale datasets"

Fig S1. Mean protein expression value to NA proportion scatter plot of NCI60(A) and CRC65(B) datasets.

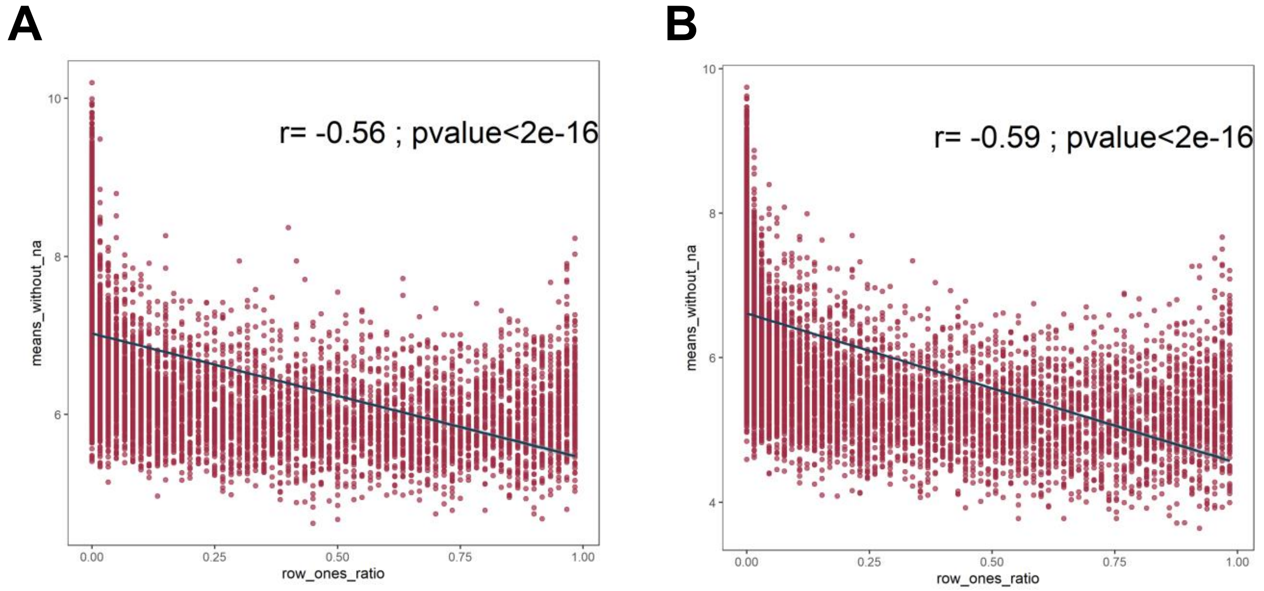


Fig S2. Low NA proportion proteins have high expression values in individual cell lines. (A) HeLa; (B) HCT-116; (C) A549; (D) NALM-6

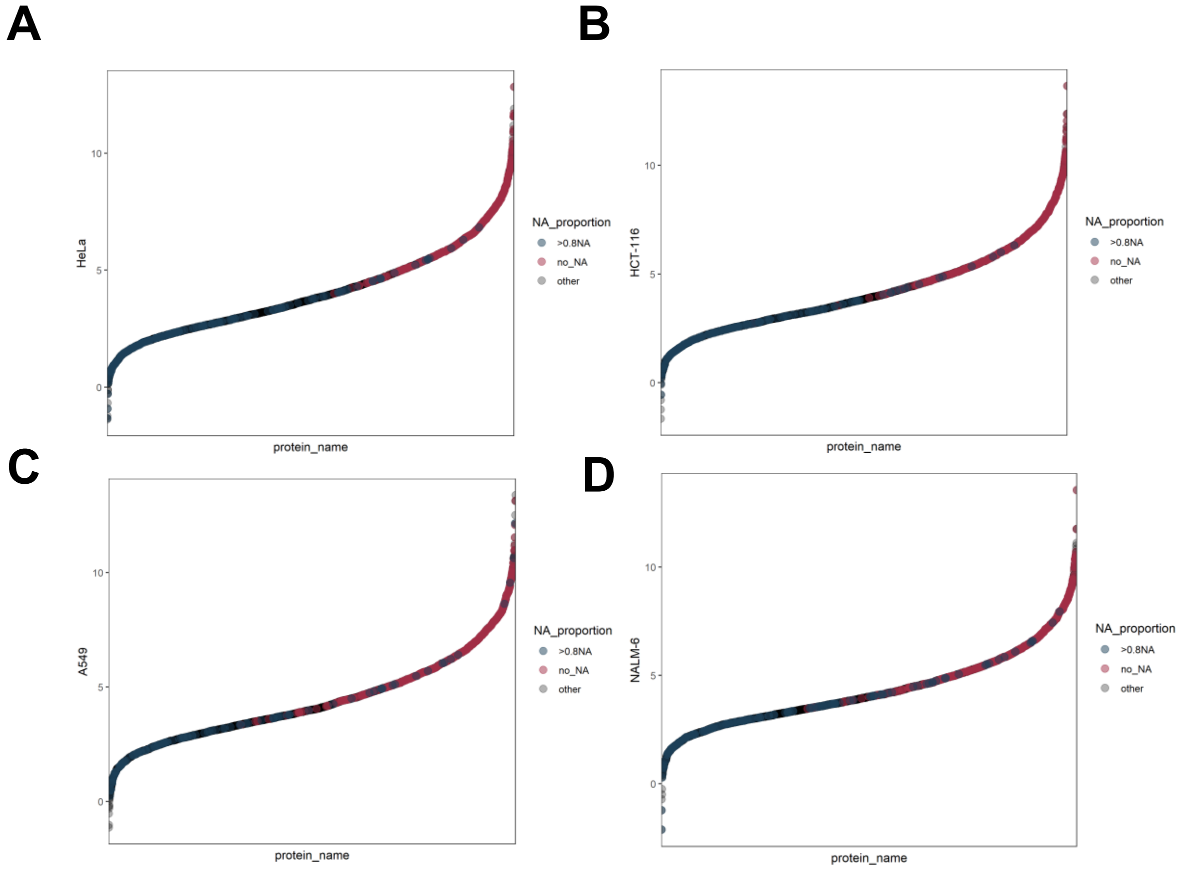

Fig S3. Sample numbers of each tissue in CCLE949 dataset.

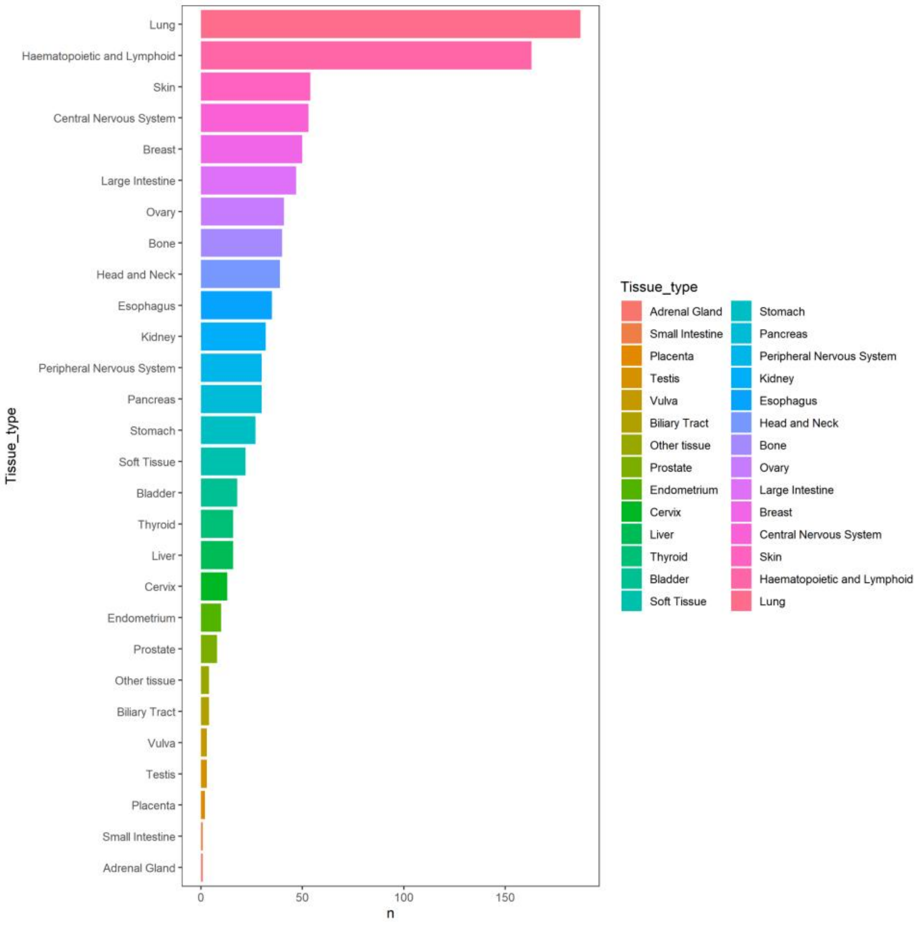


Fig S4. NA classification ROC curve of different k in the simulation dataset.

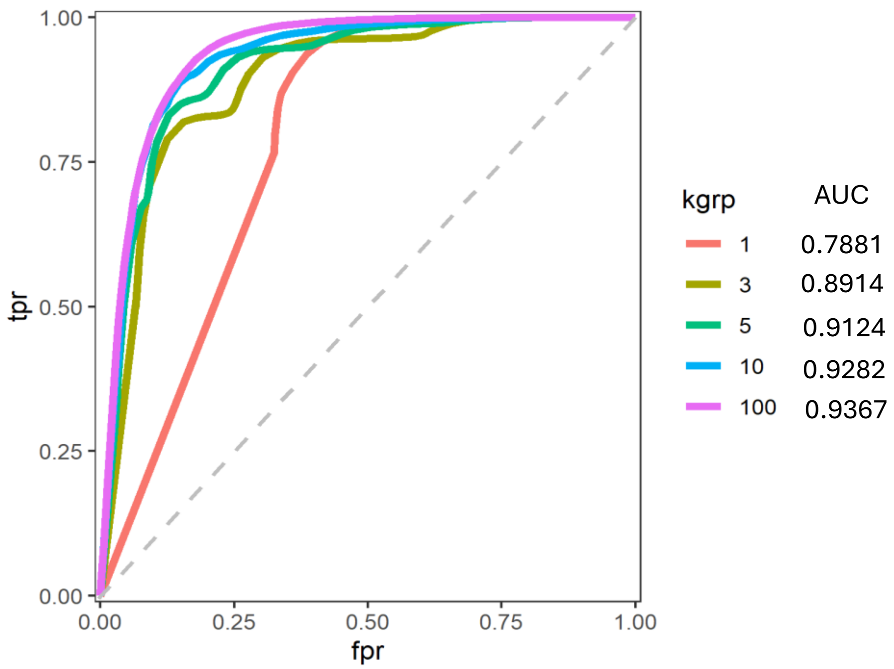

Fig S5. Hematopoietic cancer specific markers (A) and specific absence (B) found by BIND in CCLE949 dataset.

**A**

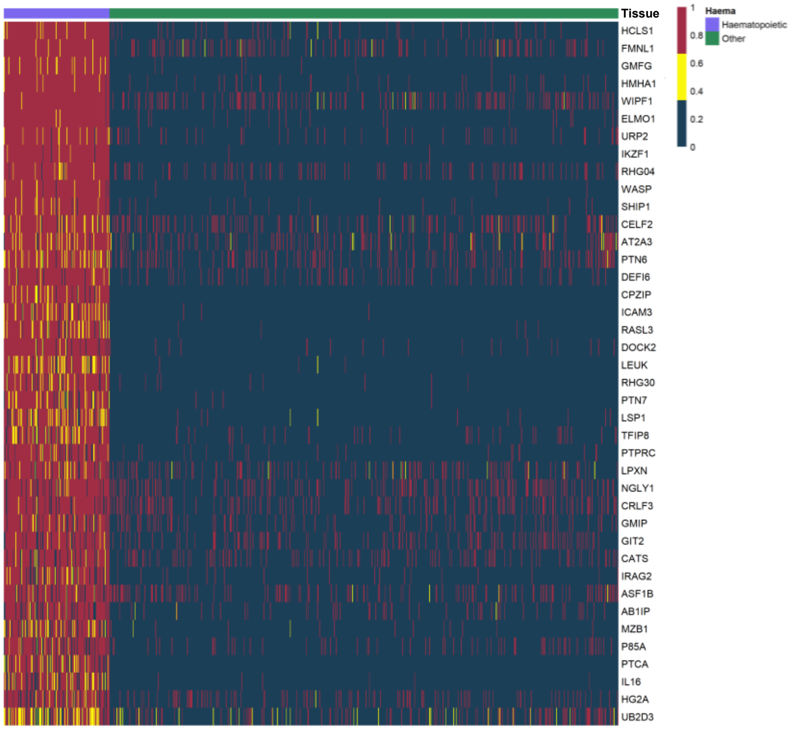


**B**

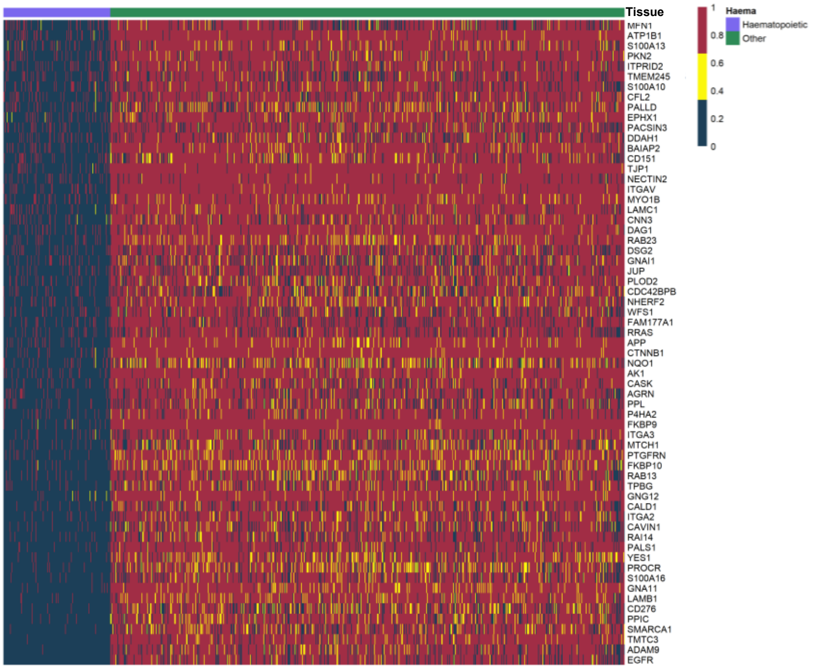

Fig S6. By improving on Spearman ρ, ρBIND rises (correlates more strongly) over Spearman ρ in a higher proportion of protein pairs with PPIs than without PPIs

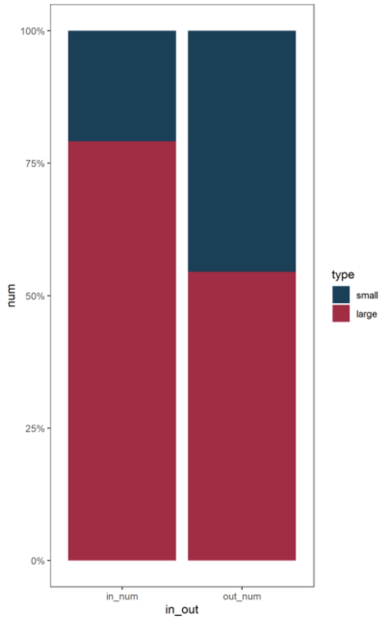


Fig S7. Fold change of ρBIND differences within and between the 20 protein complexes. Complex data were obtained from the CORUM database by selecting the top 20 complexes with the highest number of proteins in the target dataset.

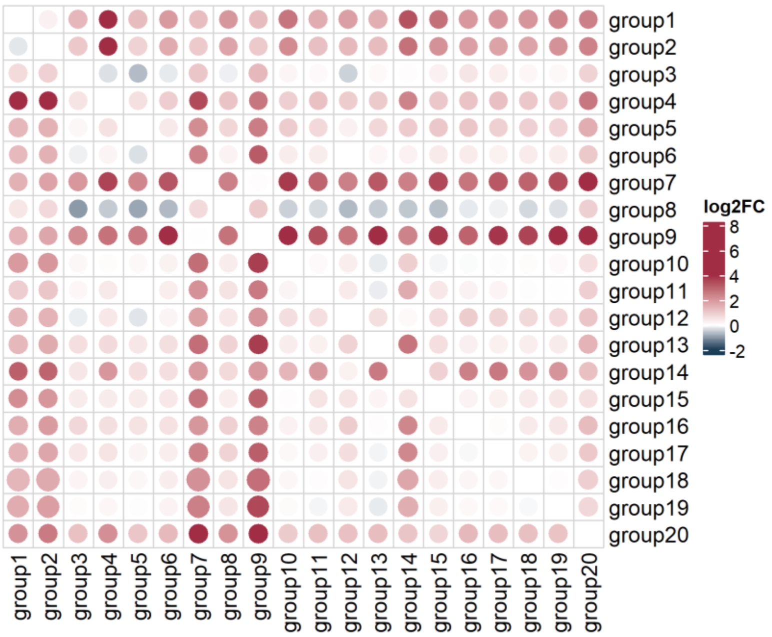

Fig S8. GO enrichment analysis of jurkat cell line markers found by BIND web server.

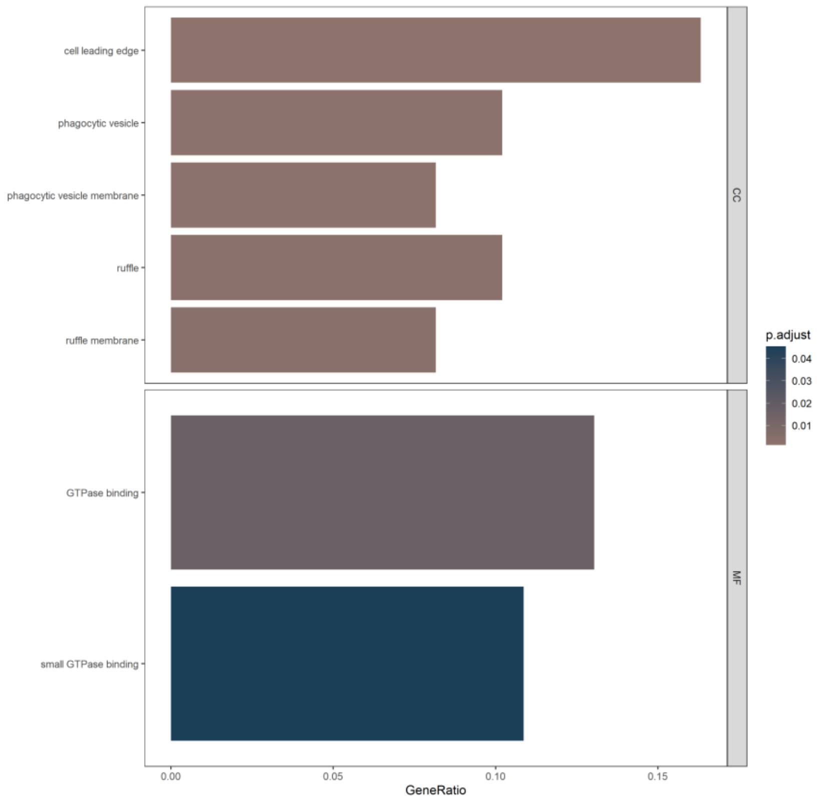
